# Galaxy-ME: A Web-based Software Resource for Interactive Analysis of Multiplex Tissue Imaging Datasets

**DOI:** 10.1101/2022.08.18.504436

**Authors:** Allison L. Creason, Cameron Watson, Qiang Gu, Daniel Persson, Luke Sargent, Yu-An Chen, Jia-Ren Lin, Shamilene Sivagnanam, Florian Wünnemann, Ajit J. Nirmal, Koei Chin, Heidi S. Feiler, Hannah Holly, Lisa M. Coussens, Denis Schapiro, Björn Grüning, Peter K. Sorger, Artem Sokolov, Jeremy Goecks

## Abstract

Highly multiplexed tissue imaging (MTI) are powerful spatial proteomics technologies that enable *in situ* single-cell characterization of tissues. However, analysis and visualization of MTI datasets remains challenging, and we developed the Galaxy-ME software hub to address this challenge. Galaxy-ME is a web-based, interactive software hub that enables end-to-end analysis and visualization of MTI datasets and is accessible to everyone. To demonstrate its utility, Galaxy-ME was used to analyze datasets obtained from multiple MTI assays in both normal and cancerous tissues. Galaxy-ME is a publicly available web resource.

## Introduction

Highly multiplexed tissue imaging (MTI) technologies, such as cyclic immunofluorescence (CycIF)^1^, multiplex immunohistochemistry (mIHC)^2^, Co-Detection by Indexing (CODEX)^3^, imaging mass cytometry (IMC)^4^, and multiplex ion beam imaging (MIBI)^5^, are powerful *in situ* spatial proteomics technologies for characterizing tissues at single-cell, and potentially subcellular, resolution. MTI has been rapidly adopted in both basic and translational research. For example, MTI has enabled identification of differences in healthy and diseased tissue organization^3^ and quantification of compositional and spatial features associated with clinical outcomes, including disease development^6^, patient survival^7–9^, and response to therapy^10^. MTI is also used extensively in several large tissue atlas consortia to create detailed 2D tissue maps for interrogating cellular and spatial relationships. Atlas consortia using MTI include the Human Cell Atlas (HCA)^11^, the Human BioMolecular Atlas Program (HuBMAP)^12^, and the Human Tumor Atlas Network (HTAN)^13^.

With the substantial growth and use of MTI technologies in the biomedical research community, there is a need for robust analytical tools and visualizations of MTI datasets. Typical MTI datasets are often tens or hundreds of gigabytes in size and include images of the tissue sample assayed with each marker protein measured, yielding a stack of 30-100 images and tens or hundreds of thousands of cells analyzed per sample. A complete software analysis workflow for MTI often uses dozens of analysis tools to perform two broad tasks: (1) primary image processing to produce single-cell feature tables with marker intensity levels, morphological information, and spatial coordinates and (2) single-cell analysis to classify individual cells and quantify spatial relationships amongst cells.

Many software applications for MTI analysis and visualization have been developed, including assay-specific computational workflows (CycIF^14^, CODEX^15^, mIHC^2,16^, and IMC^17–19^), general purpose image analysis platforms (CellProfiler^20^, QuPath^21^, and Fiji^22^), and web-based image visualization tools (Vitessce^23^ and Visinity^24^). While innovative and powerful, software for MTI analysis and visualization has several key limitations. First, most software does not support comprehensive, end-to-end analysis and visualization but instead are dependent on external software applications at different stages of image processing, data analysis, or data formatting. Software for MTI analysis and visualization that is comprehensive requires use of a command line text interface and significant computational expertise. Second, desktop applications that provide a user-friendly interface for MTI dataset analyses cannot easily be deployed or scaled across different computational infrastructure such as cloud computing platforms or computing clusters. These challenges in software tool integration, accessibility, and scalability make it difficult to analyze MTI datasets.

To address these challenges, we have developed Galaxy-MCMICRO Ecosystem (Galaxy-ME), a highly scalable web-based software hub for interactive analysis of MTI datasets that includes a graphical user interface (Fig. 1). Galaxy-ME provides a web-based software workbench for MTI analyses that is accessible to all scientists, a comprehensive tool and visualization suite for analysis of MTI datasets, and infrastructure to ensure that all analyses are scalable and reproducible (Fig. 1a). Galaxy-ME overcomes common challenges of analyzing and visualizing MTI datasets by providing end-to-end analysis workflows that reduces reliance on external software, a web-based user-friendly interface to enhance accessibility, and flexible deployment across a variety of computing infrastructures to leverage advanced computing resources. Galaxy-ME provides a total of 23 software analysis tools for (1) primary image processing to produce single-cell feature tables that include marker intensity levels, morphological information, and spatial coordinates; (2) single-cell analysis to classify individual cells and quantify spatial relationships amongst cells; and (3) interactive visualization of images and analysis results (Fig. 1b; Table 1).

**Table 1.**
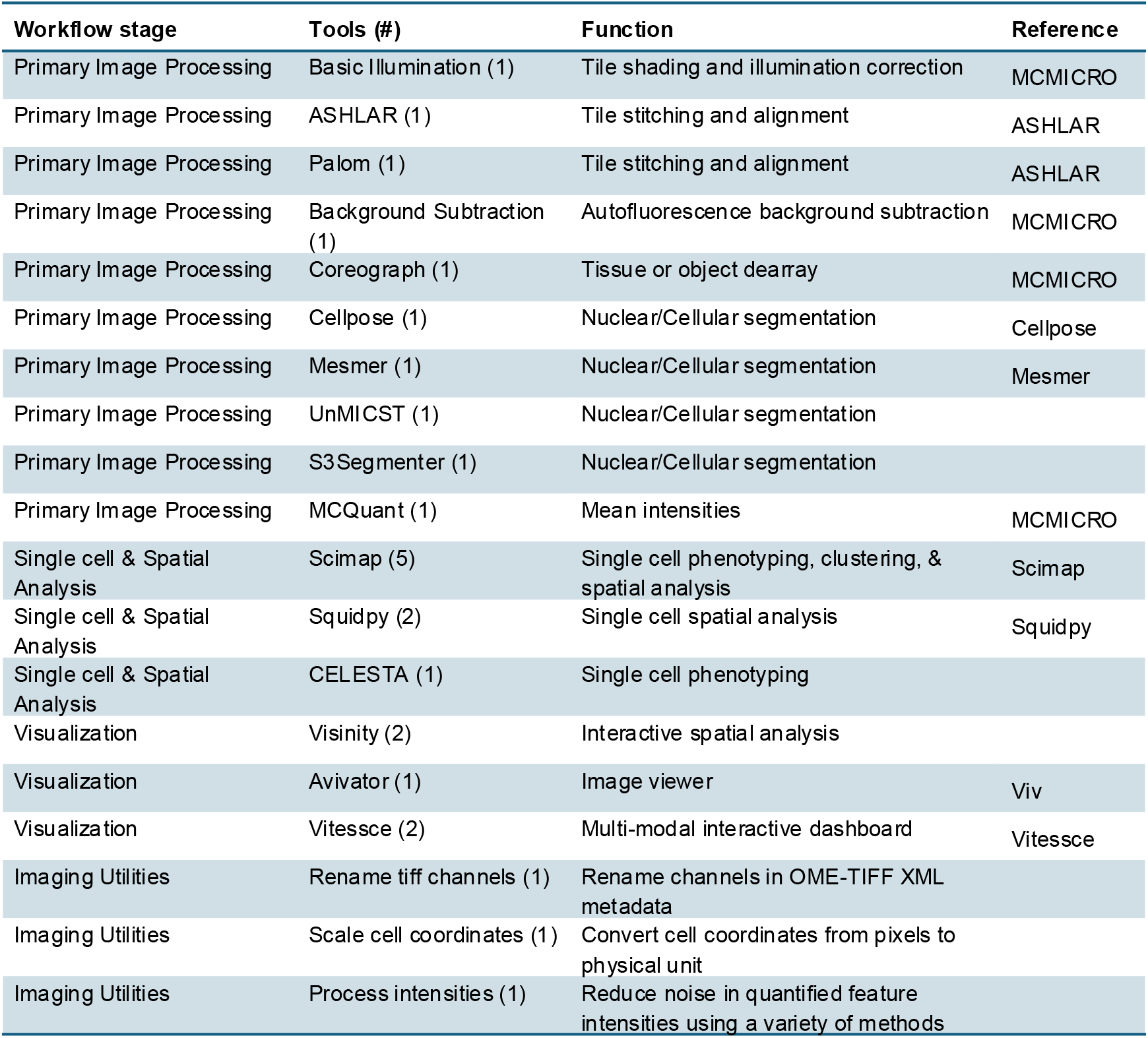
A list of all Galaxy-ME Tools organized by analysis stage. The number in parentheses represents the number of Galaxy-ME tools associated with each tool name or software package.

**Figure 1.**
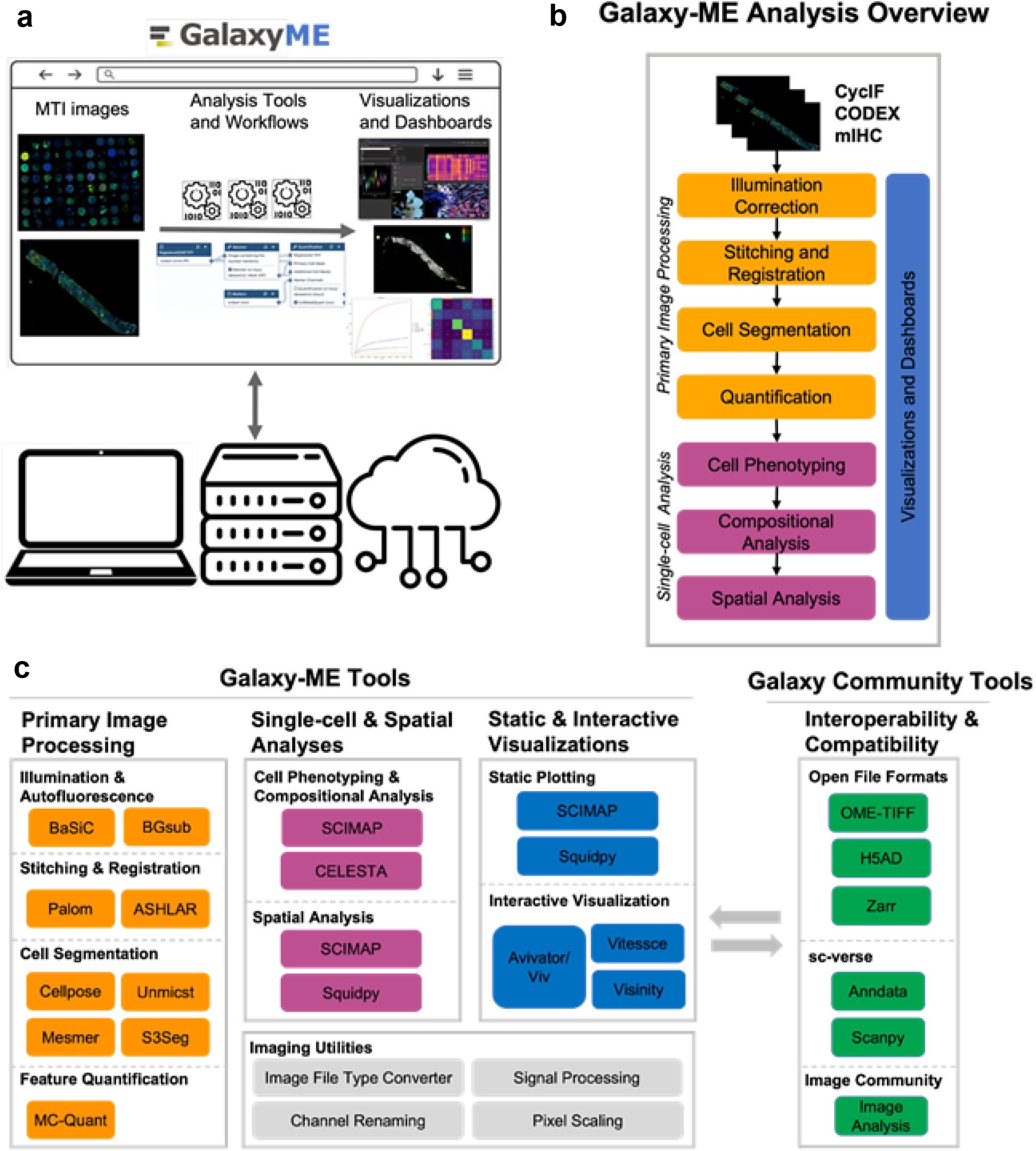
Galaxy-ME overview, tools, and visualizations. **a**) Galaxy-ME provides a web-based user interface to perform comprehensive analysis and visualization tasks for multiplexed tissue imaging datasets. Galaxy-ME can be run on a laptop, computing cluster, or a cloud computing platform. All analyses are completely reproducible, and running Galaxy-ME on a computing cluster or cloud computing platform makes it possible to complete large-scale analyses using large-scale computing resources. **b**) Overview of analysis steps and visualizations for multiplexed tissue imaging datasets available in Galaxy-ME. These include primary image processing (orange), single-cell and spatial analyses (magenta), and both interactive and static visualization options (blue). **c left)** Galaxy-ME tools organized by categories established in panel b. All tools are also listed in Table 1 with citations and descriptions of tool functions. Also shown are Galaxy-ME imaging utilities that provide convenience functions for image processing workflows; these utilities are not part of previously other software packages. **c right)** Galaxy-ME tools are interoperable and compatible with other MTI data analysis software, single-cell analysis software, and existing Galaxy tools and through use of open and standard file formats.

## Design and implementation

Galaxy-ME provides software tools and workflows for comprehensive, end-to-end analysis of MTI datasets. Galaxy-ME is built on the Galaxy computational workbench (https://galaxyproject.org/)^25,26^, an open-source platform for graphical, reproducible, and collaborative biomedical data analyses. The software and visualization tools in Galaxy-ME use best-practice analysis approaches, integrating tools from multiple atlas consortia including HCA, HuBMAP, and HTAN. All tools are open-source and have been previously validated and documented in publications, GitHub repositories, and dedicated websites (Table 1; Supplemental Table 1). Galaxy-ME tools follow community standards defined for file formats following the MITI^27^ and Bio-Formats^28^ for imaging data and sc-verse^29^ for single cell data, ensuring interoperability and extensibility.

The Galaxy-ME tools are accessible from two public web services: https://cancer.usegalaxy.org/ supports analytical tools within the cancer research community and https://spatialomics.usegalaxy.eu/ serves the European Galaxy community. If dataset or user privacy prevents use of these public web services, Galaxy-ME can also be downloaded and run locally on an institutional computer or computing cluster so that data is not uploaded or shared. Code repositories for Galaxy-ME tools are hosted on GitHub, while tool versioning and distribution is managed via the Galaxy ToolShed^30^.

Galaxy-ME tool development and publication to the ToolShed adhere to the best practices laid out in the Galaxy^25^ and Planemo^31^ documentation which help ensure reproducibility and ease-of-use for Galaxy tools (https://galaxy-iuc-standards.readthedocs.io/).

### Accessible, Scalable, and Reproducible Software Tools with the Galaxy Platform

The Galaxy computational workbench is among the most popular biomedical software analysis platforms in the world and used by thousands of scientists daily. Galaxy-ME takes advantage of the accessibility, security, scalability, and reproducibility features that Galaxy offers. Both Galaxy and Galaxy-ME are open-source and freely available. Galaxy-ME uses Galaxy’s web-based graphical user interface (GUI), making analysis of MTI datasets accessible, regardless of computational expertise. The Galaxy-ME GUI enables users to move between selecting input datasets, running analysis tools/workflows, and visualizing imaging data or single cell analysis results. Tools are implemented with options for basic and advanced parameters, to enable flexible execution though the variety of image types and formats may require users to tune parameters for their use-case. All tools are extensively documented in publications, GitHub repositories, and dedicated documentation websites which can be found in Supplemental Table 1.

Galaxy-ME’s analysis tools and visualizations are orchestrated and executed on remote computing resources by the Galaxy server. By conducting MTI analyses in Galaxy, data provenance is automatically tracked and made available via Galaxy analysis histories. Galaxy-ME tools can be run individually on selected datasets or multi-step workflows can be run to execute an entire MTI analysis using Galaxy-ME tools. By using the Galaxy framework and sufficient remoting computing resources, Galaxy-ME can scale its analyses to process collections of imaging datasets that are hundreds of terabytes in size. Galaxy uses best-practice web application and database practices as well as role-based security to ensure data and user privacy. Basic Galaxy usage is extensively documented on the Galaxy Training Network website (https://training.galaxyproject.org/)^32^.

### Primary Image Processing using MCMICRO and Additional Tools

Primary image processing in Galaxy-ME includes all the MCMICRO^18^ suite of tools, in addition to supplemental tools for key steps (Fig. 1; Table 1; Supplemental Table 1). Typical image processing steps include: (1) illumination correction between microscopy tiles (with the BaSiC^33^ tool); (2) stitching of tiles into channel mosaics (ASHLAR^34^); (3) registration of channel mosaics into a multi-channel OME-TIFF pyramid (ASHLAR, PALOM); (4) nuclear and cellular segmentation (UnMICST^35^ and S3segmenter^36^, Cellpose^37^, Mesmer^38^); and (5) cell feature quantification (MCQuant). Processing tissue microarrays requires an additional step to split the whole-slide array into individual core images (Unet Coreograph), which is performed after registration. Once the core images are split, they can be treated as any whole-slide image and processed in parallel.

Nuclear and whole-cell segmentation are challenging but critical image processing steps that can vary dramatically in performance between assays and tissues. For this reason, three segmentation tools have been integrated into Galaxy-ME so that the best segmentation method can be applied to each dataset. Additionally, the Galaxy-ME implementation of Mesmer utilizes a pre-trained model that can perform nuclear or whole-cell segmentation based on max-projection of several membrane markers, and has parameters for fine-tuning the segmentation towards a specific assay. Final outputs from primary image processing are (1) a multi-channel pyramidal OME-TIFF file that includes all image channels, (2) a mask of segmented nuclei or cells, and (3) a tabular cell feature table with mean protein marker intensities, morphological features (area, eccentricity, orientation, and solidity), and spatial coordinates for each cell identified in the segmentation mask.

Galaxy-ME also includes several utilities (Supplemental Table 1) that can be optionally run as a part of an image pre-processing pipeline. For instance, MCMICRO’s pixel-level background subtraction tool can be used to subtract autofluorescence channels from channels of interest. Alternatively, Galaxy-ME includes a utility that performs background subtraction, exposure compensation, and/or signal-to-background ratio calculation on the quantified cellular features rather than at the pixel level, which is less computationally intensive and does not require duplication of image data. Other utilities include a tool for renaming image channels in the OME-TIFF metadata, a tool for converting pixel-based cellular coordinates to a physical distance (e.g. pixels to micrometers) based on the resolution of the image, and the Bioformats image conversion tool (*bfconvert*).

### Comprehensive Single-Cell Phenotyping and Spatial Analyses Tools

Single-cell analysis tools in Galaxy-ME use cell feature tables produced by primary image processing for cell phenotyping, compositional analysis, and spatial analysis (Fig. 1). Galaxy-ME uses the common anndata format (https://anndata.readthedocs.io) for storing single-cell data. Galaxy-ME includes several convenience tools for converting between tabular format and anndata h5ad format (*scimap*.*pp*.*mcmicro_to_anndata, scimap*.*hl*.*scimap_to_csv*). Anndata format additionally allows for easy integration with existing Galaxy tool suites, like SciAp^39^ which is a Galaxy tool suite that provides single-cell analysis modules such as Scanpy^39,40^ and Seurat^41^, which provide many additional downstream analysis opportunities for Galaxy-ME users.

Galaxy-ME supports single-cell phenotyping using three complementary approaches. First, cells can be gated using a biologically-driven, semi-automated hierarchical gating approach using the SciMap^42,43^ package from MCMICRO (*scimap*.*pp*.*rescale, scimap*.*tl*.*phenotype_cells*). The output of this approach is a distinct phenotype for each cell, such as *CD8+ T-cell* or *luminal neoplastic tumor cell*. Second, the Galaxy-ME implementation of CELESTA classifies cell types using an unsupervised machine learning method that uses both protein marker intensities and cell coordinates to classify cell types in a spatially-aware manner^44^. The final approach for phenotyping cells is a data-driven approach where cells are clustered using community metrics^45,46^ (e.g. Louvain, Leiden). This approach produces clusters that can then be annotated based on the markers enriched in each cluster. The implementation of Scimap in Galaxy-ME has a clustering option; however, SciAp can also be used for this application. SciAp^39^ offers a larger range of intensity normalization, scaling, and filtering that can be performed prior to clustering. Additionally, SciAp has embedding visualization tools for helping with cell-type annotation by overlaying cluster assignments or protein intensities in the embedding space. Galaxy-ME uses spatial analysis methods in SciMap and SquidPy^47^ to quantify spatial interactions, neighbor enrichment between cell types, spatial neighborhoods, and other metrics of tissue spatial organization.

### Fast, Interactive Data Exploration With Visual Analytics Tools

Galaxy-ME includes four primary interactive visualization tools of MTI datasets. Avivator is a light-weight and web-based image viewer built with the Viv^48^ library that enables viewing multi-channel OME-TIFF images hosted on a web server. With Avivator, different image channels can be selected and visualized, and it is possible to pan and zoom around images. Avivator displays are automatically generated for OME-TIFF images uploaded to Galaxy allowing users to examine images without additional tool executions. For additional data viewing, Galaxy-ME provides a tool to create Vitessce^23^ interactive dashboards that include multi-component linked visualizations, including multiplex images, labeled segmentation masks, UMAP plots, phenotype marker enrichment, and single-cell heatmaps. Finally, Galaxy-ME also provides an interface for generating and viewing a Visinity dashboard. Visinity enables viewing of multiple imaging datasets simultaneously and facilitates multi-sample analyses. Users can perform multi-sample neighborhood analyses in Visinity and capture the quantified outputs in a file for downstream analysis^24^. While Visinity has a simple user interface as a stand-alone tool, integrating the tool into Galaxy-ME connects Visinity with the many processing and analytical tools in Galaxy-ME so that a user can perform all steps of an MTI analysis within a single interface and without moving large amounts of imaging data. Additionally, Visinity is configured on Galaxy to be executed using a two-step tool invocation. The first tool processes all input data to match the input requirements of Visinity and produces a relatively small archive dataset that persists in the user’s history. This archive dataset then serves as the primary input for the second tool, a Galaxy interactive tool that launches the Visinity dashboard in a new window. This two-tool process has two major advantages: 1) A user can revisit prior dashboards at any time due to the persistence of the processed archive in the Galaxy analysis history without the need for any re-processing. This means that dashboards can be rendered with very little waiting time. 2) Users can run the archive builder an arbitrary number of times to test the effect of altering the search radius for neighborhood generation, a critical parameter that ought to be explored thoroughly and may vary depending on the biological question. Users can then view the resulting dashboards with different neighborhood sizes side-by-side.

### Comprehensive Multi-tool Workflows for End-to-End Analyses

The Galaxy Project hosts vetted, community-developed workflows for many different types of ‘omics analyses. Galaxy workflows consist of Galaxy tools that have been pre-configured to run serially by directing the outputs of tools into the input of a subsequent tool. Multiple datasets can be invoked simultaneously and Galaxy supports the ability to combine multiple workflows together through a super-workflow. For the datasets described in this manuscript, we provide a pre-configured super-workflow that executes the core processing workflows, upstream additional tools, and downstream analyses such that the entire analysis can be reproduced in a single workflow invocation. The core workflow was optimized for each of the three imaging assays: CycIF, mIHC, and CODEX by setting a limited number of parameters at run-time along with the registered OME-TIFF image as the primary input. Links to all Galaxy-ME workflows can be found in the results section. Additionally, an example Galaxy-ME workflow can be found in the Galaxy Workflow Library, an online repository for curated high-quality community-developed workflows (https://iwc.galaxyproject.org/workflow/multiplex-tissue-microarray-analysis-main/).

## Results

Galaxy-ME brings a comprehensive collection of MTI analysis and visualization tools together in a unified framework so they can be used with each other and take advantage of Galaxy’s functionality and thousands of other tools. Using Galaxy-ME, scientists can analyze datasets from several MTI assays. To demonstrate this utility, we used Galaxy-ME to perform a fully automated and comprehensive analysis of both healthy and diseased tissue datasets from three MTI assays—CycIF, mIHC, and CODEX.

All analysis figures were generated with Galaxy-ME, and all corresponding datasets and results are available as Galaxy histories as listed in the supplementary webpage at https://github.com/goeckslab/tools-mti/blob/main/README.md. Each history is uniquely named and has a short-hand (i.e. tonsil 1, tonsil 2, crc 1, etc.) for reference in the manuscript that can be found in the supplementary web-page. The core workflow used to process datasets used in this manuscript is also available (https://cancer.usegalaxy.org/u/watsocam/w/galaxymeassayworkflow062024). A tutorial highlighting an example analysis using Galaxy-ME is available at https://training.galaxyproject.org/training-material/topics/imaging/tutorials/multiplex-tissue-imaging-TMA/tutorial.html to help get started with MTI analyses using Galaxy-ME.

### Cross-Assay Analysis of Healthy Tonsil Reveals Detailed Immune Cell Spatial Organization

Using Galaxy-ME, we quantitatively characterized the compositional and spatial landscape of a healthy human tonsil tissue resection dataset generated by the HTAN^13^ consortium where tissue sections were profiled with CycIF, mIHC, and CODEX (Fig. 2a). The healthy tonsil sample was procured as part of the Human Tumor Atlas Network (HTAN) SARDANA Trans-Network Project (Supplemental Table 2) with tissue processing and imaging of the sample as previously described^18^. Whole-slide images of the tonsil sample were acquired using mIHC and CycIF, and were cropped to match the ROI imaged using CODEX across non-adjacent sections prior to processing in Galaxy-ME.

**Figure 2.**
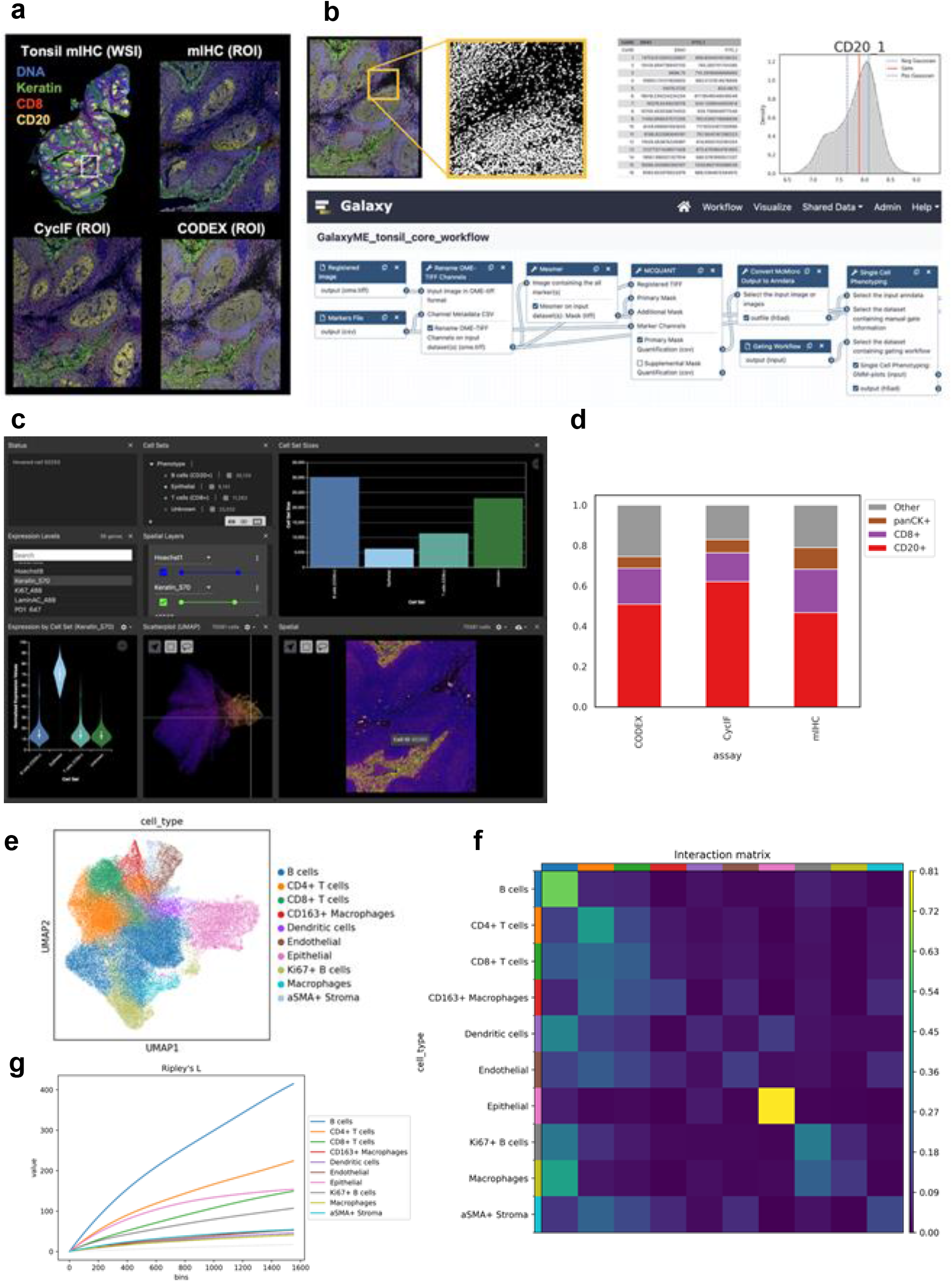
Galaxy-ME analysis of a healthy tonsil specimen using three multiplex tissue imaging assays. Galaxy-ME analysis of a healthy tonsil specimen using three multiplex tissue imaging assays. **a**) Overview of the whole slide tonsil image profiled with mIHC (top left) and a region of interest (ROI) using mIHC, CycIF, and CODEX. Images are pseudo-colored with four shared markers for DNA (blue), Keratin/PanCK (green), CD8 (red), and CD20 (yellow). Images were rendered using Avivator, the default image viewer for OME-TIFF images in Galaxy-ME. **b)** Sample outputs of Galaxy-ME’s core image processing workflow (top) and the workflow visualized in the Galaxy workflow editor (bottom). A registered OME-TIFF image is the primary input for the workflow. The workflow performs nuclear segmentation, feature quantification, and automated GMM-based cell phenotyping to create a single-cell matrix of all cells in the image. **c**) a Vitessce dashboard can be created and viewed in Galaxy-ME with no data transfer or formatting. The interactive dashboard includes views of the phenotype-labeled segmentation mask overlaid onto the registered image, compositional bar plots, UMAP representations, violin plots of marker expression, and heatmaps. Results can be filtered across visualizations, and individual cells can be highlighted using a cursor across visualizations. **d**) Stacked bar plots show cellular proportions in the tonsil ROI obtained from Galaxy-ME across each assay. Cellular phenotypes are based on three shared markers across assays: CD20+ (red), CD8+ (purple), panCK+ (brown), or Other (gray). The plot was generated using the Scimap stacked bar plot tool in Galaxy-ME. **e**) UMAP representation of cells from the CycIF tonsil image, run on a subset of 13 lineage markers generated using the Galaxy-ME Scanpy tool suite. The UMAP plot is colored by cell type, which were determined through annotation of Leiden clusters based on the enrichment of certain lineage markers. **f**) A row-normalized spatial interaction matrix using CycIF cell type annotations in panel e generated using the Galaxy-ME Squidpy tool suite. Matrix was created from a spatial neighborhood graph using a 130 micrometer radius to capture larger-scale spatial structures. Rows denote cell types, and the column colors map to the corresponding row color indicative of cell type. Epithelial cells and B-cells have a high proportion of interactions with cells of the same type. All immune cell types have relatively high proportions of interactions with B-cells. **g**) A Ripley’s L plot using CycIF cell type annotations and generated using the Galaxy-ME Squidpy tool suite. Ripley’s L is a univariate spatial metric describing the degree of clustering or dispersion exhibited by each cell type in the sample. The metric is calculated across a range of neighborhood radii, which is denoted in pixel units across the x-axis, and compared against a random point process (gray line). All cell types show strong, non-random clustering indicative of healthy tissue.

A key feature of Galaxy-ME is the ability to develop robust, standardized workflows that are generalizable across multiplex tissue imaging assays while still allowing for flexible customization. We demonstrated this functionality by building a core Galaxy-ME workflow to process datasets from all three MTI assays (Fig. 2b; Extended Data Fig. 1), including nuclear segmentation, feature quantification, and automated cell phenotyping, with custom runtime parameters to optimize platform specific differences. Platform specific customization were applied by adding pre- and post-processing tools, including pixel-level autofluorescence background subtraction for assays with fluorescence-based reporters (Extended Data Fig. 1). Execution of the core processing workflow (described above) on the three assay ROI images resulted in separate Galaxy histories corresponding to each assay (Histories: Tonsil 1, 2, 3) with the final output being a cell feature table annotated with cell phenotypes based on the expression of three shared markers (pan-cytokeratin for epithelial, CD8 for cytotoxic T-cells, and CD20 for B-cells). The resulting images, nuclear masks, and derived single-cell data were used to create Vitessce^23^ dashboards for interactive downstream analyses (Fig. 2c).

Galaxy-ME has built-in support for multi-sample analysis allowing direct comparison of common MTI outputs, such as mean cell counts based on segmented objects or cell type composition. Using this multi-sample functionality, all three assay datasets were analyzed with a combined-assay workflow (Extended Data Fig. 2). Results for each assay analyses are captured in separate Galaxy histories along with Vitessce dashboards for interactive viewing. These cell feature tables were then copied to a new Galaxy history (History: Tonsil 4) for a combined multi-modal analysis. The three assay anndata files were concatenated using anndata manipulation tools in the Galaxy Scanpy tool suite. Subsequently, static bar plots were generated using the Scimap plotting tool (sm.pl.stacked_barplot) in Galaxy to compare cell-type compositions between the three assays in both absolute and relative abundance. The mean cell count across the three assays was 71,168 cells with a standard deviation of 2,229 cells (Extended Data Fig. 3). Total cell count differences across assays are expected due to the physical separation of non-adjacent slides. Multi-sample cell type analysis using shared markers across the three assays (CD20, CD8, pan-cytokeratin) also showed concordance in broad cell-type composition (Fig. 2d, Extended Data Fig. 3).

Deeper profiling of the CycIF image (History: Tonsil 5) highlights how to apply common spatial analysis techniques using multiple popular tools with high interoperability in Galaxy-ME. The Scanpy tool suite in Galaxy was used to scale the intensities of lineage markers, generate a neighborhood graph, run UMAP dimensional reduction, and assign cells to Leiden clusters (resolution = 1.5). Leiden clusters were then annotated as cell-types based on the cluster-wide expression of lineage markers, ultimately resulting in 10 discrete cell types. A spatial neighborhood graph was constructed using the Galaxy implementation of Squidpy (*squidpy*.*gr*.*spatial_neighbors*) with a radius of 130 micrometers (200 pixels). To explore the spatial arrangement of cell types in the tonsil tissue, Squidpy was used to calculate a row-normalized interaction matrix (*squidpy*.*gr*.*interaction_matrix*) and to compute Ripley’s L statistic (*squidpy*.*gr*.*ripley*) across a range of radii from 0-1000 micrometers. Finally, spatial scatter plots were generated using Squidpy to show cell-type and marker intensity distributions across the tissue. To demonstrate all current cellular phenotyping options available in Galaxy-ME, we also used the Galaxy-ME implementation of CELESTA to assign cell types to the CycIF tonsil dataset (Extended Data Fig. 4) (History: Tonsil 6).

In-depth analysis of the CycIF image shows that Galaxy-ME can identify highly resolved immune cell types and quantify local and global cell spatial arrangements (Fig. 2e, Extended Data Fig. 4,5). Epithelial cells had a high degree of self-interaction, while immune cells interacted frequently with other immune cell types indicative of tonsillar crypt and lymphoid follicle morphology respectively (Fig. 2f). B-cells had a high degree of self-interaction and spatial clustering, and spatial scatterplots show highly localized areas of Ki67 intensity which is consistent with elevated B-cell proliferation in germinal centers (Fig. 2f,g, Extended Data Fig. 5).

### Multi-faceted Spatial Analysis of a Colorectal Cancer Provides Insight into Cellular Neighborhoods

We next applied Galaxy-ME to a HTAN colorectal cancer (CRC) resection with adjacent serial sections profiled using CycIF and mIHC^49^. Using the CRC dataset, we demonstrate two distinct approaches for multi-sample spatial analysis using Galaxy-ME: 1) a recurrent cellular neighborhood analysis on the mIHC ROI images using the Scimap implementation of spatial Latent Dirichlet Allocation (LDA); and 2) an interactive neighborhood similarity search across the CycIF ROI images using Visinity.

The CRC sample was procured as part of the Human Tumor Atlas Network (HTAN) SARDANA Trans-Network Project (Supplemental Table 2). Tissue processing and imaging of the sample has been previously described^49^. Pathologists have previously annotated this specimen with histological features used as prognostic indicators for CRC. Across the adjacent sections, seven regions of interest (ROIs) were previously defined to capture diverse histologies and tissue compositions (Extended Data Fig. 6). A whole-slide image of the CRC sample was acquired using CycIF, and was cropped to match the 7 ROI images captured using mIHC on an adjacent section prior to processing in Galaxy-ME. Registered mIHC and CycIF ROI OME-TIFF images from the colorectal cancer resection were uploaded into separate Galaxy histories (Histories: CRC 1, 2). All primary processing of the ROI images happened in parallel, as a tool can be invoked at once across all elements in a list collection in Galaxy. Primary processing was the same for the CRC datasets as the tonsil datasets. In brief, nuclei were segmented from the registered images using Mesmer, and MCQuant was used to extract single-cell features. Feature quantification files were converted to AnnData format for downstream analysis.

To characterize the spatial architecture shared across the seven ROIs, we performed an analysis of the mIHC CRC datasets to identify recurrent cellular neighborhoods (*neighborhood motifs*)^42^. Individual mIHC ROI images were concatenated into a single AnnData h5ad object with 9 markers automatically gated using Scimap’s GMM-based gating tool. Cell types were determined hierarchically resulting in 10 unique cell types across the mIHC ROI images. Stacked bar plots were generated using Scimap to visualize the cell type composition of each ROI. Cellular coordinates were converted from pixel units into micrometers using the cell coordinates scaling utility in Galaxy-ME (mIHC physical resolution = 0.5) Spatial LDA (scimap.tl.spatial_lda) was run on the concatenated file with a radius size of 30 micrometers and 10 latent motifs to be extracted from the training corpus. The number of motifs to use were determined empirically following previous methodologies^42,50^. Within the spatial LDA invocation, cellular neighborhoods were calculated for each ROI image independently, and then LDA was performed on neighborhoods from cells across all images resulting in 10 spatially recurrent motifs. The resulting ten neighborhood motifs captured a diversity of cell populations across the ROIs, including motif 3, 4, and 9 enriched in pan-cytokeratin+ epithelial cells and motifs 2, 5, 6, and 8 enriched with CD4+ T-cells, CD8+ T-cells, and CD20+ B-cells (Fig. 3a).

**Figure 3.**
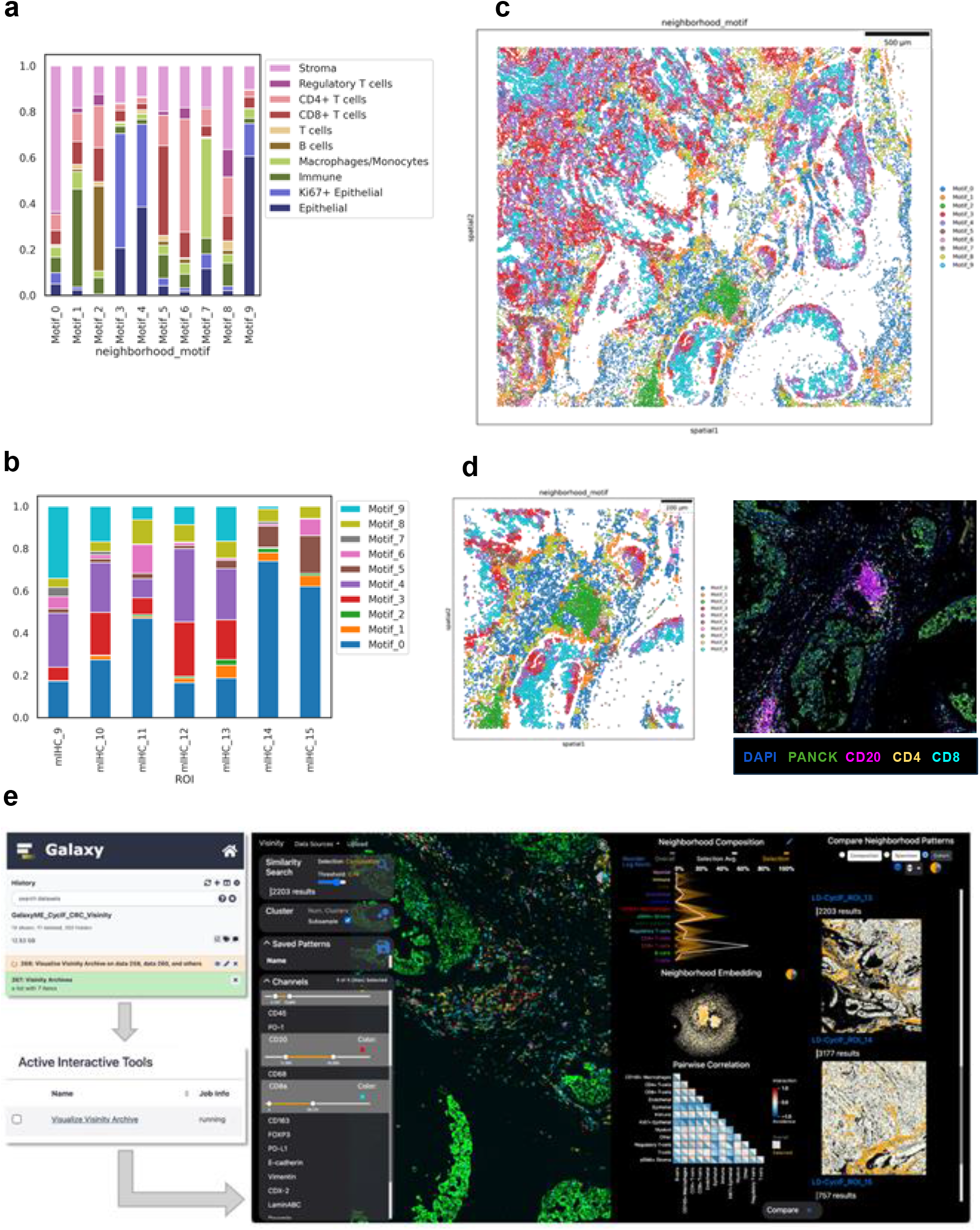
Galaxy-ME facilitates analysis of recurrent cellular neighborhoods in colorectal cancer dataset using two multiplex imaging assays. Galaxy-ME analysis results and visualizations of recurrent cellular neighborhoods in a colorectal cancer (CRC) dataset using two multiplex tissue imaging assays. Galaxy-ME results and visualizations enable deep, single-cell exploration of the CRC tumor ecosystem. **a**) Cell type composition of neighborhood motifs generated using the Scimap spatial LDA implementation in Galaxy-ME. Motifs were generated from cellular neighborhoods originating across seven CRC mIHC ROIs. **b**) Neighborhood motif composition of CRC ROIs. ROIs are ordered sequentially across the x-axis. ROIs varied substantially in neighborhood motif composition due to heterogeneous histological regions sampled. **c**) Spatial scatterplot of ROI 13 imaged using mIHC generated using Squidpy in Galaxy-ME. Cells are labeled by neighborhood motif. **d**) Cropped spatial scatterplot (left) and image viewed in Avivator (right) of an immune aggregate in ROI 13. Neighborhood Motif 2 (green) captures the immune cell diversity in this tissue region. **e**) Screenshot of Visinity archive creation and interactive tool panel in Galaxy (left), and screenshot of Visinity being used on the CycIF CRC dataset to explore recurrent spatial patterns across the cohort of ROIs (right).

Results from our analysis align well with previously published observations^49^. Neighborhood motif composition in our analysis varied across the ROIs and revealed biologically meaningful histological morphologies previously annotated (Fig. 3b). For example, ROIs 14 and 15, representing muscularis propria, were dominated by motif 0 whereas ROIs 9-13, representing invasive colonic adenocarcinoma, were predominantly composed of the pan-cytokeratin+ epithelial enriched motifs 3, 4, and 9. Our neighborhood motifs captured the various differentiation states and grades within the invasive colonic adenocarcinoma (ROIs 9-13). The poorly differentiated, solid architecture in ROI12 is composed predominantly of the Ki67+ cytokeratin+ epithelial-rich motif 3 whereas the moderately differentiated, glandular architecture of ROIs 9-11 are composed of mixed proliferative epithelial motifs 4 and 9 and contain higher proportions of the stromal rich motif 0. The mucinous pools of ROI 13 contain a balance of motifs 0,1, 3, 4, and 9 consistent with mixed moderate-high grade tumor budding into the mucinous space. Spatial plotting of the neighborhood motifs and interactive image viewing highlights an example lymphoid aggregate within the CycIF ROI 13 (Fig. 3c,d). The Scimap stacked bar plot and Squidpy spatial scatter plot tools were then used to analyze the distribution of motifs across ROIs.

As a second approach to spatial analysis, we used the Galaxy-ME interactive tool for Visinity^24^ to perform a deep exploration of the CycIF CRC datasets. The CycIF ROI images and associated downstream data, including background subtracted images, segmentation masks, and cell-type annotated cell feature tables were collected into a Galaxy history. Using the Visinity Archive building tool in Galaxy-ME, the inputs were then pre-processed for visualization and analysis (30 micrometers radius size). Once the archive builder is run, it serves as an input for the Visinity interactive tool. This two-step process with Galaxy-ME makes it simple to reuse dashboards without any dataset updates and the ability to run multiple dashboards simultaneously to find an optimal radius size for the neighborhood analysis (Fig. 3e).

Previous analysis of this CRC sample found substantial heterogeneity in the immune cell aggregates that are present in the tumor. To identify immune cell aggregates across the image cohort, we used Visinity’s built-in Cohort viewer and similarity search features. The CycIF ROI 13 was loaded into the Visinity image viewer with pseudocolor applied to the CD3, CD4, CD8, and CD20 channels. The lasso tool in Visinity can be used to highlight regions of interest and conduct a similarity search for neighborhoods with similar composition. Upon finding and outlining an immune aggregate enriched for immune markers in the ROI 13, we used the similarity search (threshold 0.75) to identify similar immune-rich tissue structures across the other ROIs. With the Cohort View in Visinity, we found CycIF ROI 14 to also be enriched for similar immune aggregates in comparison to the rest of the cohort.

Our analysis and visualization of MTI datasets from healthy human tonsil and colorectal cancer tissue specimens demonstrates the unique capabilities of Galaxy-ME to perform comparative analysis of multiple MTI samples. Galaxy-ME produces quantitative metrics for both single-cell compositional and spatial tissue organization across samples, and sample results are aggregated and visualized. With Galaxy-ME two complementary multi-sample spatial analysis approaches, Scimap LDA RCN analysis and Visinity neighborhood similarity analysis, that can be applied to a multi-sample cohort. To the best of our knowledge, Galaxy-ME is the only software platform capable of multi-sample and assay agnostic spatial analyses coupled with built-in primary image processing as well.

## Availability and Future Direction

The source code for the Galaxy-ME tools is available at https://github.com/goeckslab/tools-mti. All Galaxy-ME tools are available for use via a web browser on the Galaxy Cancer Server (https://cancer.usegalaxy.org/), the EU Galaxy server (https://spatialomics.usegalaxy.eu/), and the main U.S. Galaxy server (https://usegalaxy.org/).

Web-based analysis and visualization of multiplexed tissue imaging datasets remains a challenging problem as these large datasets require using many disparate analysis tools and visualizations together, often in an iterative and interactive manner. The Galaxy-ME software democratizes access, analysis, and visualization of MTI datasets and empowers all scientists to analyze MTI datasets regardless of their informatics expertise. Galaxy-ME enables comprehensive end-to-end analysis of MTI datasets—including image processing, spatial analysis, and interactive visualization—using a graphical web-based user interface. Building this expansive tool suite on Galaxy, an open-source biomedical computational workbench used daily across the world, brings many benefits. By leveraging the Galaxy workbench, the entire Galaxy-ME tool suite is accessible, reproducible, secure, and scalable. Further, Galaxy-ME users can readily use Galaxy’s fully-featured built-in workflow engine, data management, and privacy capabilities with individual user accounts. Galaxy-ME tools can be connected with more than 10,000 other tools available via the Galaxy Tool Shed^30^, to create analyses that extend far beyond imaging, such as using machine learning tools^51^ or integration with other omics data analysis workflows for multimodal analysis.

There are limitations of building Galaxy-ME on the Galaxy platform, which is a general-purpose graphical biomedical workbench. One important limitation is that the Galaxy user interface cannot be easily adapted for MTI analyses because it must provide a consistent design for the thousands of tools integrated into Galaxy. This limitation is especially relevant because MTI analyses often benefit from a highly visual user interface with dynamic controls that provide immediate visual feedback about results of analysis steps. We have mitigated this limitation in Galaxy-ME by including the powerful visualization tools Visinity and Vitessce to provide a rich interactive experience for MTI analyses. Galaxy-ME offers insights about how future open-source MTI analysis systems can balance the tradeoff between general-purpose interfaces for biomedical data analyses and specialized highly interactive interfaces for MTI. While Galaxy’s consistent user interface has drawbacks for MTI analyses, the consistent interface is beneficial to the Galaxy-ME user experience in many ways as it provides a consistent and unified interface to all Galaxy-ME’s tools. Another limitation is that Galaxy-ME’s graphical user interface may be slower than code-based approaches for large bulk analyses.

However, Galaxy has a Python-based library for Galaxy tools and workflows that can be used to automate large analyses^52^. Automated analysis results can be viewed in Galaxy’s graphical interface or they can be downloaded and used outside of Galaxy. Future work with Galaxy-ME is likely to address the need to simultaneously and interactively perform cell segmentation, phenotyping, and spatial analytics tasks. Interactive analyses across all MTI analysis steps will help researchers optimize and understand their analysis results. There are also opportunities to add additional tools and visualizations that extend Galaxy-ME beyond 2D single-cell analysis to focus on quantification of larger units of biological organization such as functional tissue units^36^ and 3D tissue organization as well as analyses of smaller, subcellular characteristics of cells. Finally, integration of additional artificial intelligence (AI) and machine learning (ML) tools for automated processing of MTI datasets is likely to be valuable in the future. AI/ML tools may be useful for identifying similar tissue regions from MTI datasets and for using features of MTI datasets to predict phenotypic or clinical outcomes.

## Data Availability

The tonsil image datasets analyzed are available at https://www.synapse.org/MCMICRO_images, and the CRC images are available at https://www.synapse.org/#!Synapse:syn47164089. In addition, all datasets will be made available through the HTAN Data Portal. The list of Galaxy histories used in this work is available at https://github.com/goeckslab/tools-mti/blob/main/README.md.

## Code Availability

The Galaxy-ME code repository, including galaxy tool wrappers and Docker files, are fully open-source and available at: https://github.com/goeckslab/tools-mti. A tutorial demonstrating an example analysis using Galaxy-ME is available at: https://training.galaxyproject.org/training-material/topics/imaging/tutorials/multiplex-tissue-imaging-TMA/tutorial.html.

## Acknowledgements

This project was carried out with major support from the OHSU SMMART Program, National Cancer Institute (NCI) grants U24CA284167 and U24CA231877, and NCI Human Tumor Atlas Network (HTAN) Research Centers at OHSU (U2CCA233280) and Harvard Medical School (U2CCA233262), and Prospect Creek Foundation. D.S. and F.W. are supported by the German Federal Ministry of Education and Research (BMBF 01ZZ2004) as part of the HiGHmed consortium.

## Figures and Tables

**Supplemental Table 1:** Documentation and code repositories for Galaxy-ME tools. Galaxy-ME tools with links to documentation and the developer’s code repositories.

**Supplemental Table 2:** HTAN Biospecimen IDs and File IDs for accession of the SARDANA CRC images. The tonsil and CRC datasets used in this manuscript can be found using the HTAN data portal, and accessed using the identifiers provided in the table.

**Extended Data Figure 1:** Galaxy-ME background subtraction workflow addition and tool execution. **a**) Example of how Galaxy workflows can be customized to suit a specific assay’s needs. For processing the CyCIF datasets, a background subtraction tool was inserted prior to the core processing workflow. **b**) Screenshot of the Galaxy-ME background subtraction tool and the parameters that can be set. Advanced options are shown but are hidden by default. All settings have best-practice default values and are hidden initially for ease of use. **c**) A background-subtracted CycIF Tonsil image viewed in Avivator via Galaxy-ME. Avivator dashboards are immediately available when an OME-TIFF dataset is uploaded or generated in Galaxy.

**Extended Data Figure 2:** Multi-sample analysis using Galaxy-ME super workflows. Screenshot of the Galaxy workflow editor showing the super-workflow that can reproduce all processing and analysis on the tonsil dataset on all three imaging modalities, in addition to the extended analysis of the CycIF tonsil image. All images are processed using the core processing workflow, which is included as a step in the super-workflow. Image outputs can be viewed in a Vitessce dashboard, automatically generated for each modality by the super-workflow.

**Extended Data Figure 3:** Composition and cell phenotyping of the tonsil datasets. **a**) Absolute abundance of broad cell classifications across all three imaging modalities for the tonsil datasets. **b**) UMAP plots for the CycIF tonsil dataset colored by lineage markers, created using the Galaxy SCIAP tool suite. These plots aid in unsupervised cluster-based cell phenotype assignment.

**Extended Data Figure 4:** Example of spatially-aware cell type calling on the CycIF tonsil dataset using Galaxy-ME implementation of CELESTA. **a**) Output of expression probability plotting tool in the Galaxy-ME CELESTA tool for two markers, keratin and aSMA. These aid in setting probability thresholds for cell type calling algorithm in CELESTA. **b**) Final cell types called using CELSTA on the CycIF tonsil dataset.

**Extended Data Figure 5:** Spatial scatterplots of tonsil Ki67 mean cellular intensity and cell types from CycIF extended analysis. **a**) Spatial scatterplot created using the Galaxy-ME squidpy spatial scatterplot tool, colored by Ki67 mean intensity. **b**) Spatial scatterplot created using the Galaxy-ME squidpy spatial scatterplot tool, colored by all cell types. **c**) Spatial scatterplot created using the Galaxy-ME squidpy spatial scatterplot tool, colored by just B-cells and Ki67+ B-cells. The tool has the functionality to filter which cell types to plot. Ki67+ B-cells correspond with the spatial scatterplot of Ki67 intensities in panel **a**, demonstrating how spatial scatterplots can be used to validate cellular phenotyping.

**Extended Data Figure 6:** WSI placement and phenotypic composition of CycIF and mIHC CRC ROIs. **a**) ROI bounding boxes for the CRC mIHC (left) and CycIF (right) images. **b**) Cell phenotype relative abundance across CRC ROIs for both CycIF and mIHC.

## References

1. Lin, J.-R. et al. Highly multiplexed immunofluorescence imaging of human tissues and tumors using t-CyCIF and conventional optical microscopes. Elife 7, (2018).

2. Tsujikawa, T. et al. Quantitative Multiplex Immunohistochemistry Reveals Myeloid-Inflamed Tumor-Immune Complexity Associated with Poor Prognosis. Cell Rep. 19, 203–217 (2017).

3. Goltsev, Y. et al. Deep Profiling of Mouse Splenic Architecture with CODEX Multiplexed Imaging. Cell 174, 968–981.e15 (2018).

4. Giesen, C. et al. Highly multiplexed imaging of tumor tissues with subcellular resolution by mass cytometry. Nat. Methods 11, 417–422 (2014).

5. Angelo, M. et al. Multiplexed ion beam imaging of human breast tumors. Nat. Med. 20, 436–442 (2014).

6. Risom, T. et al. Transition to invasive breast cancer is associated with progressive changes in the structure and composition of tumor stroma. Cell 185, 299–310.e18 (2022).

7. Liudahl, S. M. et al. Leukocyte Heterogeneity in Pancreatic Ductal Adenocarcinoma: Phenotypic and Spatial Features Associated with Clinical Outcome. Cancer Discov. 11, 2014–2031 (2021).

8. Keren, L. et al. A Structured Tumor-Immune Microenvironment in Triple Negative Breast Cancer Revealed by Multiplexed Ion Beam Imaging. Cell 174, 1373–1387.e19 (2018).

9. Jackson, H. W. et al. The single-cell pathology landscape of breast cancer. Nature 578, 615–620 (2020).

10. Berry, S. et al. Analysis of multispectral imaging with the AstroPath platform informs efficacy of PD-1 blockade. Science 372, (2021).

11. Regev, A. et al. The Human Cell Atlas. Elife 6, (2017).

12. HuBMAP Consortium. The human body at cellular resolution: the NIH Human Biomolecular Atlas Program. Nature 574, 187–192 (2019).

13. Rozenblatt-Rosen, O. et al. The Human Tumor Atlas Network: Charting tumor transitions across space and time at single-cell resolution. Cell 181, 236–249 (2020).

14. Eng, J. et al. cmIF: A Python Library for Scalable Multiplex Imaging Pipelines. in Mathematical and Computational Oncology 37–43 (Springer International Publishing, 2019).

15. Schürch, C. M. et al. Coordinated Cellular Neighborhoods Orchestrate Antitumoral Immunity at the Colorectal Cancer Invasive Front. Cell 183, 838 (2020).

16. Blom, S. et al. Systems pathology by multiplexed immunohistochemistry and whole-slide digital image analysis. Sci. Rep. 7, 15580 (2017).

17. Schapiro, D. et al. histoCAT: analysis of cell phenotypes and interactions in multiplex image cytometry data. Nat. Methods 14, 873–876 (2017).

18. Schapiro, D. et al. MCMICRO: a scalable, modular image-processing pipeline for multiplexed tissue imaging. Nat. Methods 19, 311–315 (2022).

19. Czech, E., Aksoy, B. A., Aksoy, P. & Hammerbacher, J. Cytokit: a single-cell analysis toolkit for high dimensional fluorescent microscopy imaging. BMC Bioinformatics 20, 448 (2019).

20. McQuin, C. et al. CellProfiler 3.0: Next-generation image processing for biology. PLoS Biol. 16, e2005970 (2018).

21. Bankhead, P. et al. QuPath: Open source software for digital pathology image analysis. Sci. Rep. 7, 16878 (2017).

22. Schindelin, J. et al. Fiji: an open-source platform for biological-image analysis. Nat. Methods 9, 676–682 (2012).

23. Keller, M. S. et al. Vitessce: integrative visualization of multimodal and spatially resolved single-cell data. Nat. Methods 22, 63–67 (2025).

24. Warchol, S. et al. Visinity: Visual spatial neighborhood analysis for multiplexed tissue imaging data. IEEE Trans. Vis. Comput. Graph. 29, 106–116 (2023).

25. Galaxy Community. The Galaxy platform for accessible, reproducible, and collaborative data analyses: 2024 update. Nucleic Acids Res. 52, W83–W94 (2024).

26. Goecks, J., Nekrutenko, A., Taylor, J. & Galaxy Team. Galaxy: a comprehensive approach for supporting accessible, reproducible, and transparent computational research in the life sciences. Genome Biol. 11, R86 (2010).

27. Schapiro, D. et al. MITI minimum information guidelines for highly multiplexed tissue images. Nat. Methods 19, 262–267 (2022).

28. Linkert, M. et al. Metadata matters: access to image data in the real world. J. Cell Biol. 189, 777–782 (2010).

29. Virshup, I. et al. The scverse project provides a computational ecosystem for single-cell omics data analysis. Nat. Biotechnol. 41, 604–606 (2023).

30. Blankenberg, D. et al. Dissemination of scientific software with Galaxy ToolShed. Genome Biol. 15, 403 (2014).

31. Bray, S. et al. The Planemo toolkit for developing, deploying, and executing scientific data analyses in Galaxy and beyond. Genome Res. 33, 261–268 (2023).

32. Hiltemann, S. et al. Galaxy Training: A powerful framework for teaching! PLoS Comput. Biol. 19, e1010752 (2023).

33. Peng, T. et al. A BaSiC tool for background and shading correction of optical microscopy images. Nat. Commun. 8, 14836 (2017).

34. Muhlich, J. L. et al. Stitching and registering highly multiplexed whole-slide images of tissues and tumors using ASHLAR. Bioinformatics 38, 4613–4621 (2022).

35. Yapp, C. et al. UnMICST: Deep learning with real augmentation for robust segmentation of highly multiplexed images of human tissues. Commun Biol 5, 1263 (2022).

36. Saka, S. K. et al. Immuno-SABER enables highly multiplexed and amplified protein imaging in tissues. Nat. Biotechnol. 37, 1080–1090 (2019).

37. Stringer, C., Wang, T., Michaelos, M. & Pachitariu, M. Cellpose: a generalist algorithm for cellular segmentation. Nat. Methods 18, 100–106 (2021).

38. Greenwald, N. F. et al. Whole-cell segmentation of tissue images with human-level performance using large-scale data annotation and deep learning. Nat. Biotechnol. 40, 555–565 (2022).

39. Moreno, P. et al. User-friendly, scalable tools and workflows for single-cell RNA-seq analysis. Nat. Methods 18, 327–328 (2021).

40. Wolf, F. A., Angerer, P. & Theis, F. J. SCANPY: large-scale single-cell gene expression data analysis. Genome Biol. 19, 15 (2018).

41. Satija, R., Farrell, J. A., Gennert, D., Schier, A. F. & Regev, A. Spatial reconstruction of single-cell gene expression data. Nat. Biotechnol. 33, 495–502 (2015).

42. Nirmal, A. J. et al. The Spatial Landscape of Progression and Immunoediting in Primary Melanoma at Single-Cell Resolution. Cancer Discov. 12, 1518–1541 (2022).

43. Nirmal, A. J. & Sorger, P. K. SCIMAP: A Python toolkit for integrated spatial analysis of multiplexed imaging data. arXiv [q-bio.QM] (2024).

44. Zhang, W. et al. Identification of cell types in multiplexed in situ images by combining protein expression and spatial information using CELESTA. Nat. Methods 19, 759–769 (2022).

45. Traag, V. A., Waltman, L. & van Eck, N. J. From Louvain to Leiden: guaranteeing well-connected communities. Sci. Rep. 9, 5233 (2019).

46. Levine, J. H. et al. Data-driven phenotypic dissection of AML reveals progenitor-like cells that correlate with prognosis. Cell 162, 184–197 (2015).

47. Palla, G. et al. Squidpy: a scalable framework for spatial omics analysis. Nat. Methods 19, 171–178 (2022).

48. Manz, T. et al. Viv: multiscale visualization of high-resolution multiplexed bioimaging data on the web. Nat. Methods 19, 515–516 (2022).

49. Lin, J.-R. et al. Multiplexed 3D atlas of state transitions and immune interaction in colorectal cancer. Cell 186, 363–381.e19 (2023).

50. Chen, Z., Soifer, I., Hilton, H., Keren, L. & Jojic, V. Modeling multiplexed images with spatial-LDA reveals novel tissue microenvironments. J. Comput. Biol. 27, 1204–1218 (2020).

51. Gu, Q. et al. Galaxy-ML: An accessible, reproducible, and scalable machine learning toolkit for biomedicine. PLoS Comput. Biol. 17, e1009014 (2021).

52. Sloggett, C., Goonasekera, N. & Afgan, E. BioBlend: automating pipeline analyses within Galaxy and CloudMan. Bioinformatics 29, 1685–1686 (2013).

