## Supplemental Information for "Galaxy-ME: A Web-based Software Resource for Interactive Analysis of Multiplex Tissue Imaging Datasets"

**Supplemental Table 1.** Documentation and code repositories for Galaxy-ME tools

| Tool | Documentation | Code repository |
|---|---|---|
| Basic Illumination | https://mcmicro.org/ | https://github.com/labsyspharm/basic-illumination |
| ASHLAR | https://mcmicro.org/ | https://github.com/labsyspharm/ashlar |
| Palom | https://github.com/labsyspharm/palom | https://github.com/labsyspharm/palom |
| Background Subtraction | https://mcmicro.org/ | https://github.com/SchapiroLabor/Background_subtraction |
| Coreograph | https://mcmicro.org/ | https://github.com/HMS-IDAC/UNetCoreograph |
| Cellpose | https://www.cellpose.org/ | https://github.com/MouseLand/cellpose |
| Mesmer | https://deepcell.readthedocs.io/en/master/ | https://github.com/vanvalenlab/deepcell-tf |
| UnMICST | https://mcmicro.org/ | https://github.com/HMS-IDAC/UnMicst |
| S3Segmenter | https://mcmicro.org/ | https://github.com/HMS-IDAC/S3segmenter |
| MCQuant | https://mcmicro.org/ | https://github.com/labsyspharm/quantification |
| Scimap | https://scimap-doc.readthedocs.io/en/latest/ | https://github.com/labsyspharm/scimap |
| Squidpy | https://squidpy.readthedocs.io/en/stable/ | https://github.com/scverse/squidpy |
| CELESTA | https://github.com/plevritis-lab/CELESTA | https://github.com/plevritis-lab/CELESTA |
| Visinity | https://github.com/labsyspharm/visinity | https://github.com/labsyspharm/visinity |
| Avivator | http://viv.gehlenborglab.org/ | https://github.com/hms-dbmi/viv |
| Vitessce | https://vitessce.io/ | https://github.com/vitessce/vitessce |

**Supplemental Table 2.** HTAN Biospecimen IDs and File IDs for accession of the SARDANA CRC images.

| Dataset | Assay | ROI/WSI | HTAN Biospecimen ID | HTAN File ID |
|---|---|---|---|---|
| SARDANA CRC | CycIF | WSI | HTA13_1_6 | HTA13_1_7001 |
| SARDANA CRC | mIHC | ROI 09 | HTA13_1_7 | HTA13_1_9131 |
| SARDANA CRC | mIHC | ROI 10 | HTA13_1_7 | HTA13_1_9132 |
| SARDANA CRC | mIHC | ROI 11 | HTA13_1_7 | HTA13_1_9133 |
| SARDANA CRC | mIHC | ROI 12 | HTA13_1_7 | HTA13_1_9134 |
| SARDANA CRC | mIHC | ROI 13 | HTA13_1_7 | HTA13_1_9135 |
| SARDANA CRC | mIHC | ROI 14 | HTA13_1_7 | HTA13_1_9136 |
| SARDANA CRC | mIHC | ROI 15 | HTA13_1_7 | HTA13_1_9137 |

**Extended Data Figure 1.** Background subtraction workflow addition and tool execution

a

b

c

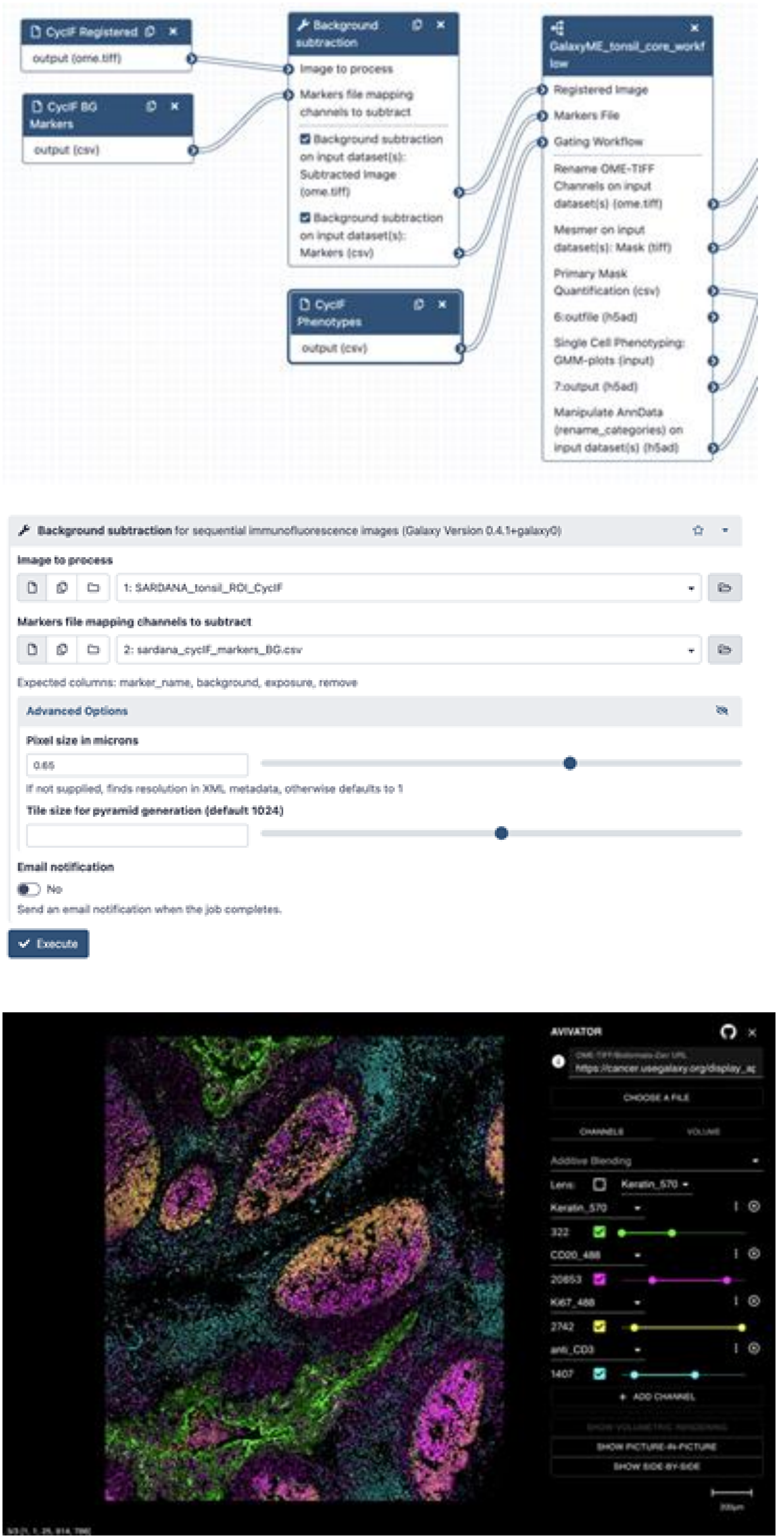

**Extended Data Figure 2.** Multi-sample analysis using Galaxy-ME super-workflows

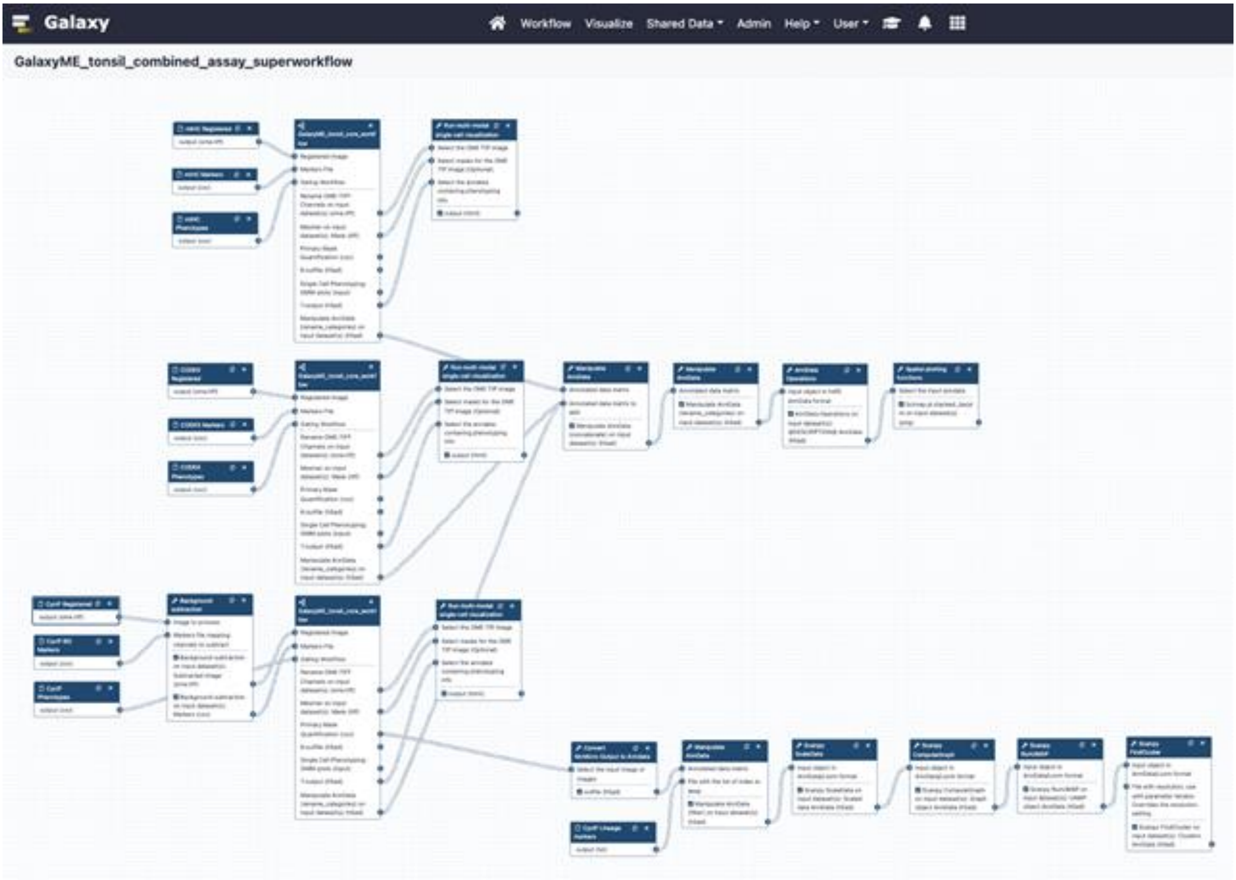

**Extended Data Figure 3.**

**a**

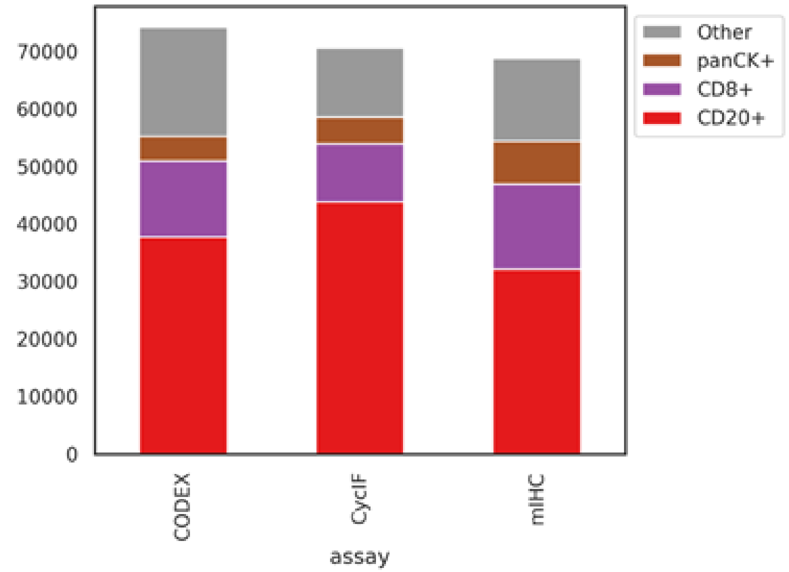


**b**

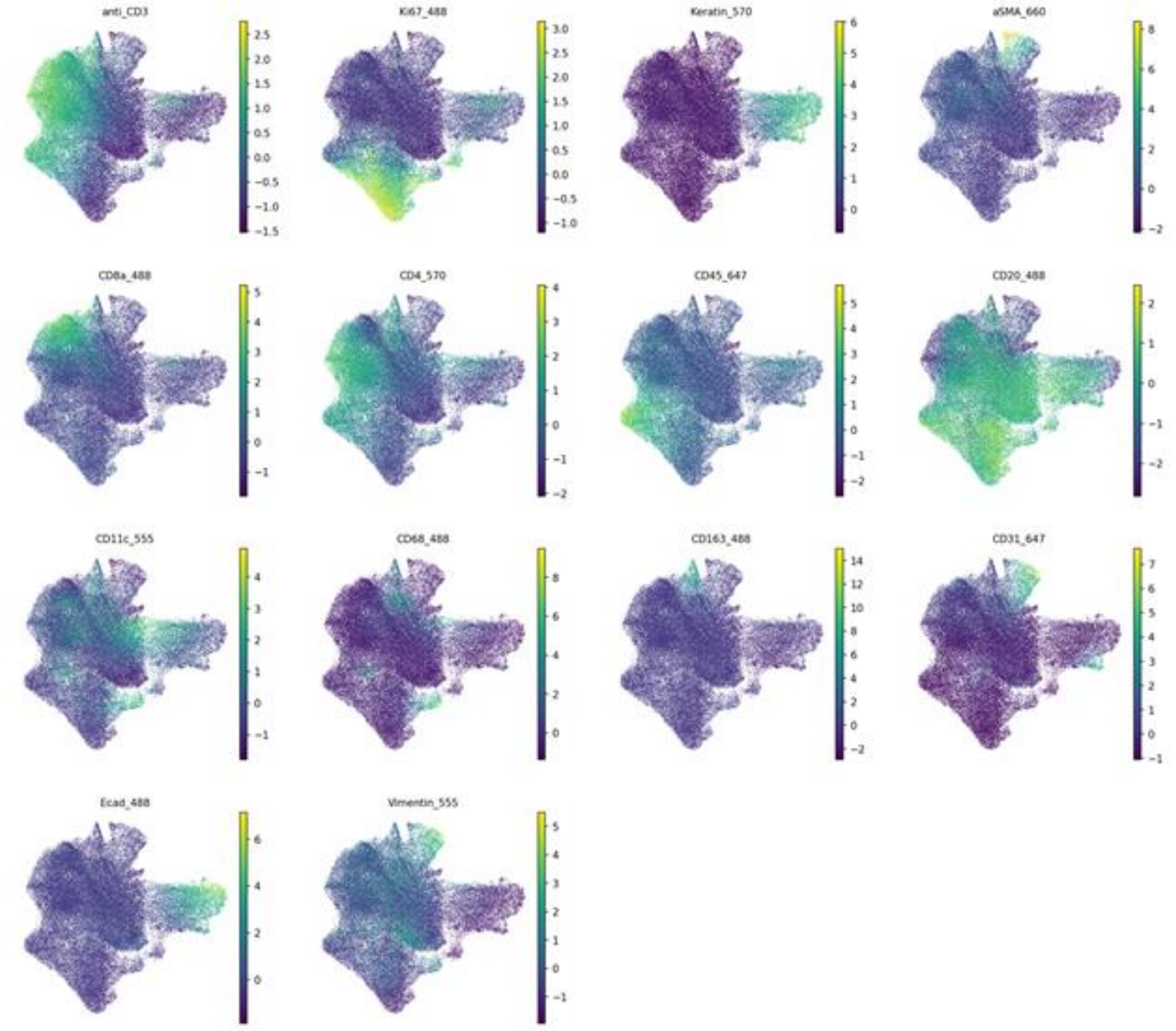

**Extended Data Figure 4.** Example of spatially-aware cell type calling on the CycIF tonsil dataset using Galaxy-ME implementation of CELESTA

**a**

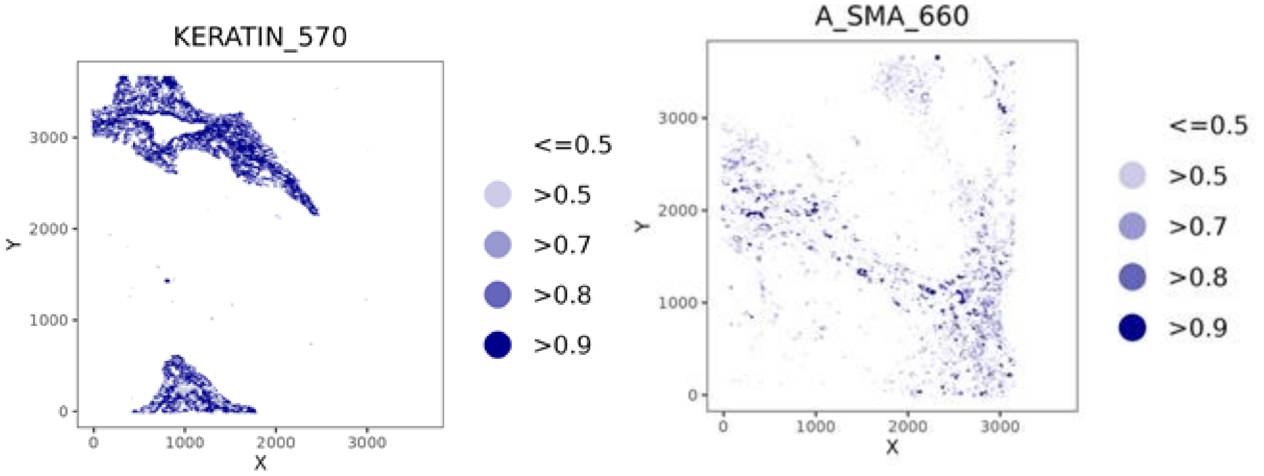


**b**

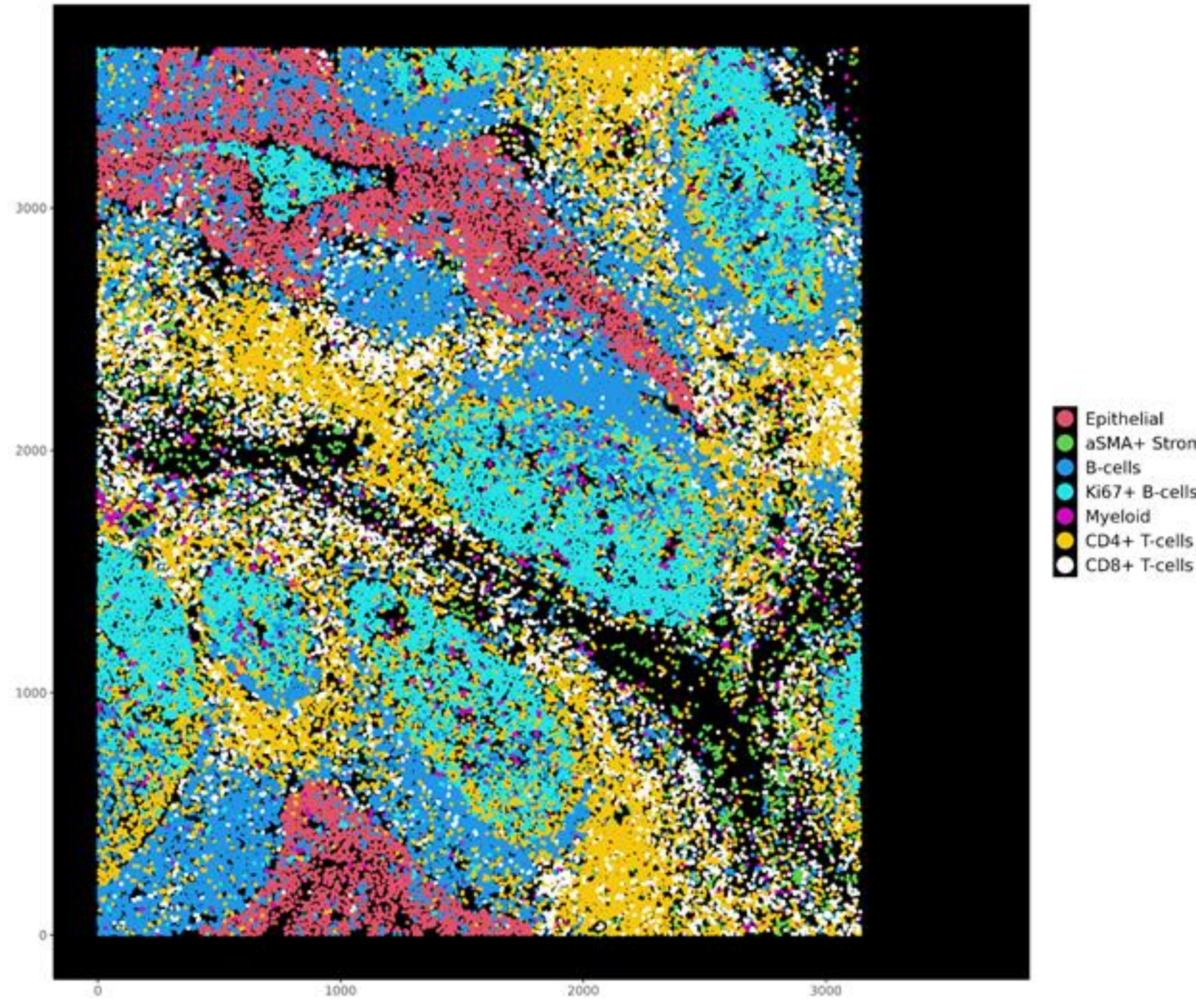

**Extended Data Figure 5.** Spatial scatterplots of tonsil Ki67 mean cellular intensity and cell types from CycIF extended analysis

a

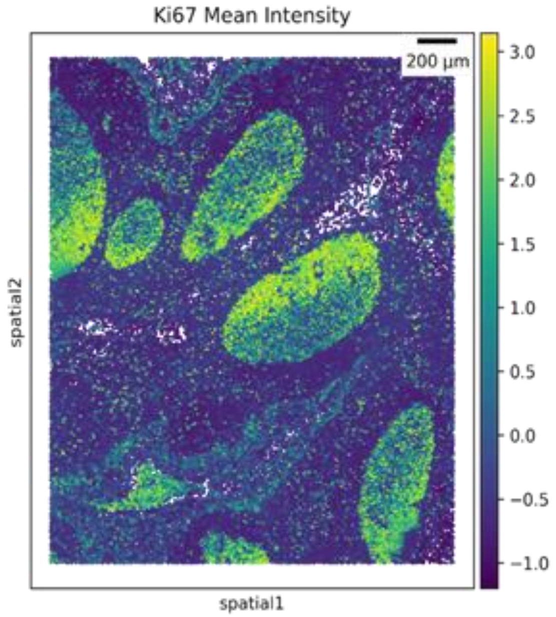


b

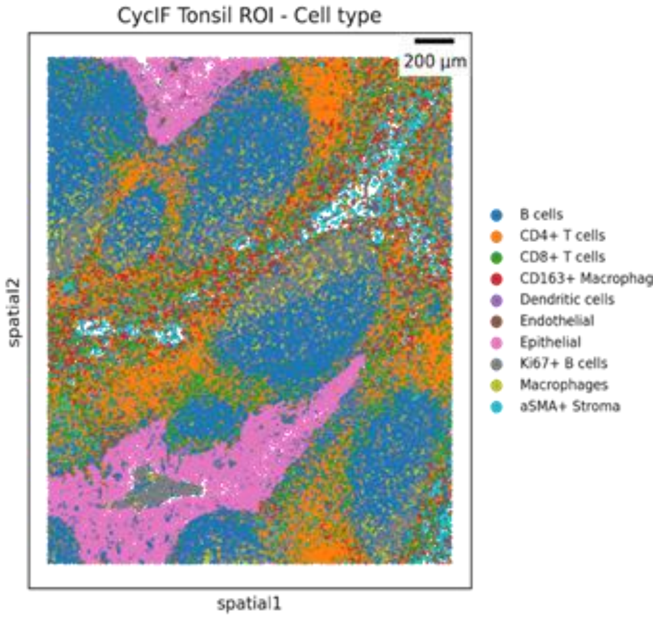


c

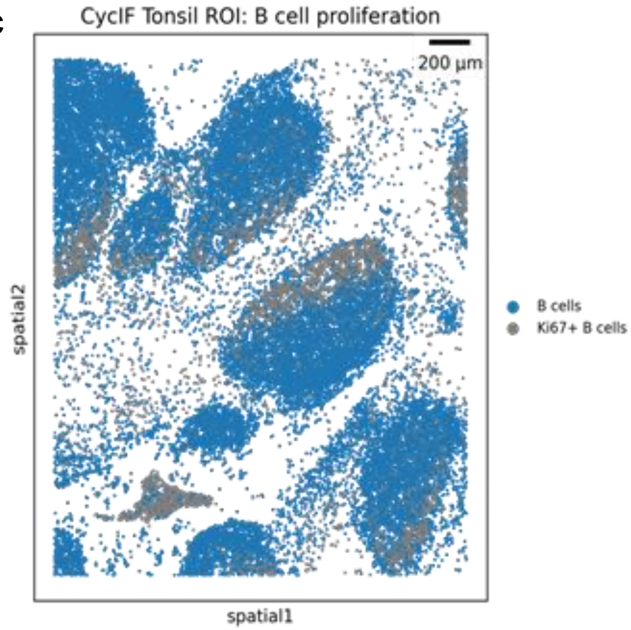

**Extended Data Figure 6.** WSI placement and phenotypic composition of CycIF and mIHC CRC ROIs

**a**

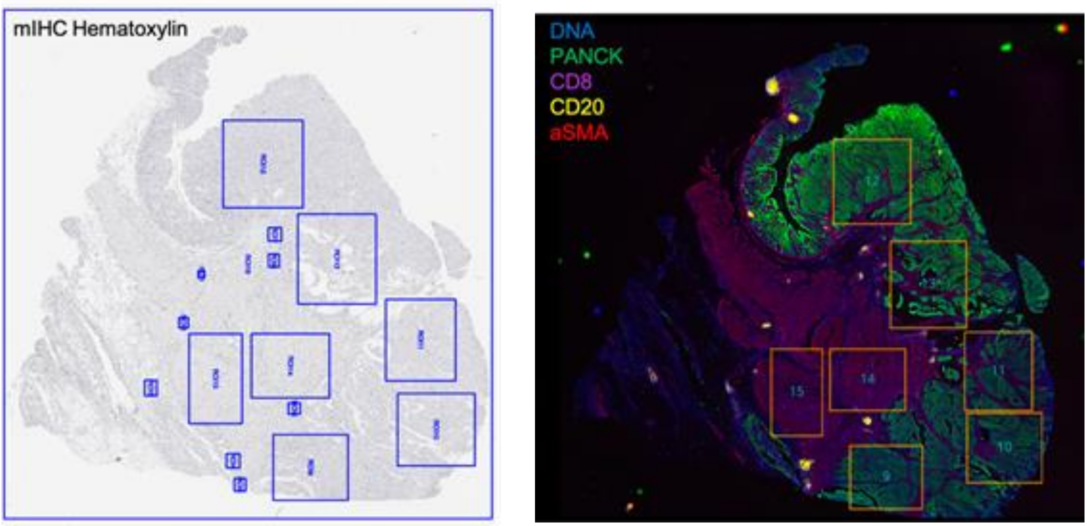


**b**

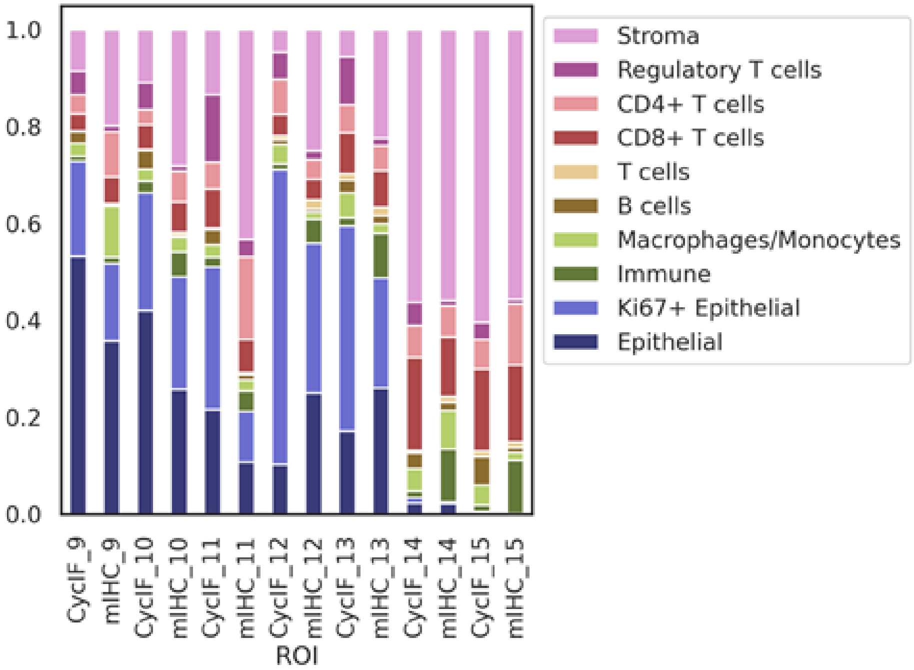
